# THE CORRELATION BETWEEN REDOX ACTIVITY AND ANTIMICROBIAL PROPERTIES OF PYOCYANIN FROM *Pseudomonas aeruginosa*

**DOI:** 10.1101/2025.02.17.638708

**Authors:** Mansi Saini, Subrata K Das, Dharmesh, Gaurav Dutt, Kishlay Kant Singh, Divya Prakash

**Author notes:** Both authors made equal contributions (Mansi Saini, Subrata K Das).

## Abstract

Pseudomonas aeruginosa, a metabolically versatile gram-negative bacterium, has garnered attention for its ability to thrive in diverse environments and its capacity to produce pyocyanin, a secondary metabolite with multifunctional properties. Pyocyanin, a redox-active phenazine compound, plays a critical role in mediating the ecological competitiveness of *P. aeruginosa* through its antimicrobial, biofilm-modulating, and reactive oxygen species (ROS)-generating activities. Beyond its contributions to bacterial virulence, pyocyanin demonstrates significant potential in various industrial and biomedical applications due to its redox properties and ability to function under diverse environmental conditions. This study investigates the electrochemical behaviour and pH-dependent antimicrobial activity of pyocyanin to evaluate its applicability in environmental and medical fields. Soil-derived *P. aeruginosa* isolates were cultured for pyocyanin (PYO) production, and the pigment was characterized using UV-visible spectroscopy to get its spectral integrity. Electrochemical analysis through cyclic voltammetry revealed enhanced redox activity in acidic environments and stable functionality in alkaline conditions. Antimicrobial assays demonstrated that pyocyanin exhibited optimal activity at neutral to slightly alkaline pH, effectively inhibiting bacterial and fungal growth, while extreme acidic conditions reduced its efficacy.

The findings highlight pyocyanin’s versatility as both a redox mediator and an antimicrobial agent. In medical contexts, its pH-sensitive activity aligns well with physiological conditions, offering promise for combating multidrug-resistant pathogens. Future optimization of pyocyanin biosynthesis, particularly through cost-effective and scalable methods, could unlock its full potential in biotechnological and therapeutic innovations.

## INTRODUCTION

*Pseudomonas aeruginosa* is a gram-negative, facultatively anaerobic, rod-shaped bacterium with a ubiquitous presence in natural and artificial environments, including soil, water, sewage, and clinical settings [1,2,3]. Owing to its remarkable metabolic versatility and environmental adaptability, *P. aeruginosa* has emerged as a significant opportunistic pathogen, particularly in immunocompromised individuals and patients with chronic illnesses, where it is implicated in severe infections such as pneumonia, urinary tract infections, and wound infections [2][4]. A critical determinant of its virulence is the production of an array of bioactive secondary metabolites, including phenazine derivatives, among which pyocyanin (PYO) is the most extensively studied [2][5].

Pyocyanin is a redox-active, nitrogen-containing heterocyclic compound with the molecular structure 5-methyl-1-hydroxyphenazine [6]. It exists as a zwitterion at physiological pH, facilitating its diffusion across biological membranes [5][7][8]. Pyocyanin exhibits distinct redox states: oxidized (blue), monovalently reduced (colorless), and divalently reduced (red), with its colour being pH-dependent—blue-green under neutral and alkaline conditions and pink-red in acidic environments [8][2]. Its biosynthesis is tightly regulated by environmental factors, particularly iron availability, with low iron concentrations enhancing its production [9][10]. Given its redox-active nature, pyocyanin engages in electron transfer reactions, playing a pivotal role in the generation of reactive oxygen species (ROS) such as superoxide and hydrogen peroxide [7]. These ROS contribute to oxidative stress-mediated cytotoxicity, conferring pyocyanin with antimicrobial properties that target both gram-positive and gram-negative bacteria, as well as fungal pathogens [6,7].

The redox potential of pyocyanin enables it to act as an electron shuttle, facilitating electron transfer between reducing agents (e.g., NADH, NADPH) and molecular oxygen, thereby promoting the formation of highly reactive superoxide anion radicals [8][11]. The resultant oxidative stress disrupts cellular homeostasis, induces lipid peroxidation, and impairs essential biomolecular functions, culminating in microbial cell death [7]. This mechanism underlies pyocyanin’s antagonistic activity against competing microbial species, reinforcing its role as a virulence factor in *P. aeruginosa* infections [12].

Beyond its role in pathogenesis, pyocyanin has garnered significant attention for its biotechnological and industrial applications [8][13,14]. Its capacity to generate ROS and facilitate electron transfer has been exploited in biocontrol strategies for agricultural applications, as well as in the development of novel antimicrobial agents. Additionally, pyocyanin functions as an effective electron mediator in microbial fuel cells (MFCs), enhancing extracellular electron transport and improving energy conversion efficiency [8][14]. Moreover, its electroactive properties have been explored in organic electronics, particularly in organic light-emitting devices (OLEDs), where it demonstrates advantageous electronic characteristics, including low voltage requirements and tunable emission spectra [15].

Despite its promising applications, challenges persist regarding the large-scale biosynthesis and economic feasibility of pyocyanin [16]. While *P. aeruginosa* remains the primary source of natural pyocyanin production, optimizing its biosynthetic pathways—particularly through the exploitation of marine-derived *P. aeruginosa* strains with enhanced yield and functional activity—may improve production scalability and cost-effectiveness [8][17,18]. Furthermore, the emergence of antimicrobial resistance necessitates the development of alternative therapeutics [19]. Given its redox-mediated antimicrobial activity, pyocyanin represents a promising candidate for combating multidrug-resistant pathogens such as Staphylococcus aureus and Candida albicans [20,21].

This study aims to elucidate the correlation between the redox properties and antimicrobial efficacy of pyocyanin, with a particular focus on its activity under varying pH conditions. A comprehensive understanding of pyocyanin’s electrochemical behaviour and ROS-generating capabilities will provide insights into its mechanism of action and its potential applications in diverse biomedical and environmental contexts, including wastewater treatment and infection control. Through this investigation, we seek to expand the scope of pyocyanin’s functional utility, particularly in addressing antimicrobial resistance and advancing biotechnological innovations.

## MATERIALS AND METHOD

### 1. Isolation and Identification of *Pseudomonas aeruginosa*

Soil samples were collected from a municipal landfill to isolate *Pseudomonas aeruginosa*, a bacterium known for its pigment production and versatile metabolic profile. Sampling was conducted at a depth of 15 cm below the surface to minimize surface contamination. The samples were transferred into sterile polyethylene bags and transported to the laboratory under aseptic conditions.

For bacterial enrichment, 1 g of soil was homogenized in 9 mL of sterile distilled water, and the suspension was vortexed to ensure uniform mixing. A 1 mL aliquot of the suspension was inoculated into 100 mL of nutrient broth within sterile conical flasks and incubated at 37°C for 24 hours under static conditions. Post-incubation, the cultures were subjected to shaking at 150 rpm in an orbital incubator shaker for an additional 24 hours to promote pigment production. The emergence of a green-blue hue in the culture medium, indicative of pyocyanin production, confirmed the presence of *Pseudomonas aeruginosa*.

### 2. UV-Visible Spectroscopic Analysis of Pyocyanin

Purified pyocyanin pigment was subjected to UV-Visible spectrophotometric analysis to ascertain its absorption characteristics and purity. The pigment was dissolved in analytical-grade hydrochloric acid (HCl) to achieve a stable solution. Spectral measurements were performed across a wavelength range of 300–700 nm using a UV-Vis spectrophotometer. The recorded absorption maxima served as a fingerprint for pyocyanin, ensuring its chemical identity and purity.

### 3. Electrochemical Characterization via Cyclic Voltammetry

The electrochemical properties of pyocyanin were investigated using cyclic voltammetry (CV) to understand its redox behavior under varying pH conditions. Solutions of pyocyanin were prepared with precise volumes of HCl or sodium hydroxide (NaOH) to achieve different pH levels (2.5 µL, 5 µL, 7.5 µL, and 10 µL of reagent additions). Electrochemical measurements were conducted on a Phadke Potentiostat electrochemical workstation (Phadke Instruments Pvt. Ltd., Mumbai, India). The resulting cyclic voltammograms provided insights into the electron transfer kinetics and potential redox-active sites of pyocyanin.

### 4. Antimicrobial Activity of Pyocyanin at Varied pH Conditions

#### 4.1 Plate Exposure Method

To cultivate environmental bacteria, including endospore formers, expose LB agar plates to the open air for one hour, allowing airborne microorganisms to settle. Subsequently, incubate the plates at 37°C overnight to promote colony formation, with endospore-forming Gram-positive bacteria like Bacillus and Clostridium expected to appear as distinct colonies; these can then be selected and picked for further analysis or culturing based on their characteristic morphologies.

#### 4.2 Preparation of pH-Adjusted Pyocyanin Solutions

The pH of pyocyanin solutions was modulated by the incremental addition of either HCl (acidic conditions) or NaOH (basic conditions). Acidic solutions were prepared by adding 2.5 µL, 5.5 µL, 6.5 µL, 7.5 µL, and 10 µL of HCl to 10 mL of pyocyanin. For basic conditions, 20 µL, 40 µL, 60 µL, and 80 µL of NaOH were added in the lowest pH sample (sample with 10 µl HCL addition) to raise the pH with varying pH levels.

#### 4.3 Antibacterial Assay

The antibacterial efficacy of pyocyanin was assessed via the agar disk diffusion method. Mixed bacterial colonies, isolated from soil samples, were cultured on nutrient agar plates prepared under sterile conditions. Using a sterile spreader, bacterial suspensions were evenly distributed across the agar surface. Sterile paper disks, impregnated with pyocyanin solutions of varying pH, were placed onto the inoculated plates. The plates were incubated at 37°C for 24 hours, and the diameter of the inhibition zones was measured to quantify antibacterial activity.

#### 4.4 Antifungal Assay

Antifungal activity was evaluated using fungal strains isolated from soil samples. The fungi were cultured on nutrient agar plates, and uniform lawns were prepared. Pyocyanin solutions adjusted to different pH levels were applied to sterile paper disks, which were then placed on the fungal culture plates. The plates were incubated at 37°C for 48 hours. Zones of inhibition around the disks were measured to determine the antifungal potential of pyocyanin.

## RESULT AND DISSICUSSION

### 1. Isolation and Identification of *Pseudomonas aeruginosa*

Soil samples, sourced from a landfill site at a depth of 15 cm, were processed under sterile conditions. A one-gram aliquot of the soil sample was suspended in 9 mL of distilled water and vortexed to achieve a homogeneous suspension. A 100 µL sample of this suspension was then plated on nutrient agar and incubated at 37°C for 24 hours. Upon incubation, colony with distinctive green-blue pigmentation emerged along with other different colonies. The green-blue colony was presumptively identified as *Pseudomonas aeruginosa*, based on their characteristic colony morphology and pigment production. To confirm this identification, we cultured the colony in nutrient broth overnight and then streaked it onto nutrient agar plate. The culture in culture tube and nutrient agar plate were continued upto 48 hrs at 37°C incubator. A green-blue pigment appeared in both liquid and solid culture. Further, we spin down the culture to separate the blue coloured supernatant and separated in two aliquot. In one aliquot, we added few drops of strong HCL, it transform into red colour (Figure 1). These observations confirmed the presence of pyocyanin, a phenazine-derived pigment secreted by the bacteria, which imparted the culture with a green-blue coloration in neutral range and red colour in acidic condition.

**Figure 1:**
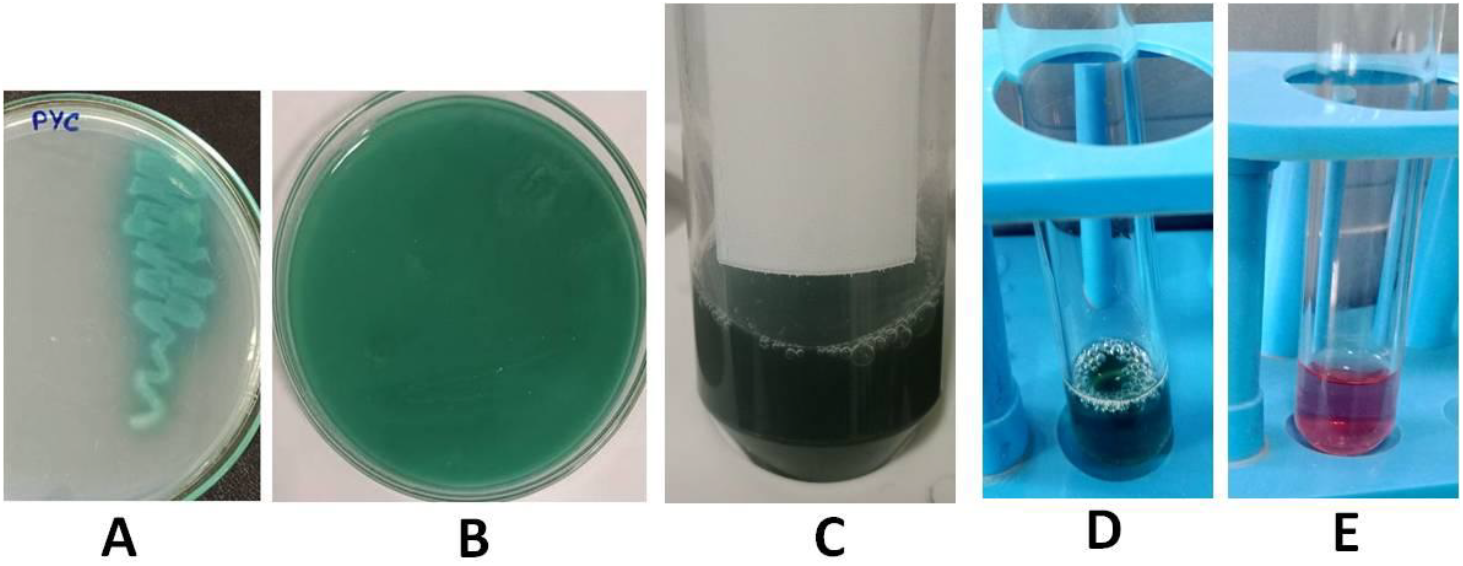
Identification of *P. aeruginosa*, (A) Growth of bacteria on nutrient agar, characterized by the secretion of blue-colored pyocyanin. (B) The entire agar plate displays bluish-green pyocyanin secretion after 72 hours of incubation. (C) A tube containing bacterial culture exhibits bluish-green pyocyanin after 72 hours. (D) The supernatant obtained from (C) post-centrifugation reveals bluish-green pyocyanin. (E) The addition of strong hydrochloric acid to the supernatant from (C) results in the formation of red-colored pyocyanin.

### 2. Redox homeostasis and survival of *Pseudomonas aeruginosa*

Upon standing for 24 hours, liquid cultures of P. aeruginosa exhibit a distinct blue ring at the air-liquid interface, which progressively expands downwards, eventually causing the entire culture to turn greenish-blue. This phenomenon is attributed to the presence of pyocyanin (PYO), a redox-active compound that acts as an extracellular electron shuttle [22]. In oxygen-limited environments, such as the bottom of the culture tube, PYO serves as an alternative electron acceptor, facilitating redox homeostasis and promoting cell survival in anaerobic niches [23]. The reduced form of PYO is re-oxidized upon diffusion to the oxygen-rich surface [23]. Thus, a green-yellow gradient observed near the air surface of standing cultures is due to the decreasing concentration of the oxidized blue form of PYO, mirroring the oxygen tension gradient. PYO primarily accepts electrons from NADH, generated during carbon source oxidation, which allows *P. aeruginosa* to survive in oxygen-poor conditions by transporting electrons to distant acceptors [23]. Furthermore, PYO stimulates pyruvate excretion, potentially supporting anaerobic survival through pyruvate fermentation. Under aerobic conditions, PYO accepts electrons from molecular oxygen, while under anaerobic conditions; *P. aeruginosa* utilizes a pyruvate fermentation pathway to sustain survival. This fermentation process regenerates NAD++, which is essential for glycolysis, allowing the bacterium to survive long-term in the anaerobic layers of biofilms and in cystic fibrosis lungs, even though it cannot sustain significant anaerobic growth [24].

When we keep standing the liquid culture in culture tube for 24 hrs, we observed that a blue ring on the upper part of liquid culture. The ring become increased towards down with increasing time and the whole culture become greenish blue in colour. Pyocyanin is auto-oxidized by oxygen, it function as an extracellular electron shuttle. The cells at the bottom are limited for oxygen but pyocyanin could serve as an alternative electron acceptor. The reduced pyocyanin become reoxidized after diffusion to the oxygen-rich surface. Thus, pyocyanin contributes to the redox homeostasis and survival of the cells in the deeper anaerobic niches [23].

Pyocyanin (PYO) is a redox-active compound that can accept and donate electrons. Its ability to easily cross biological membranes allows it to act as a mobile electron carrier for *P. aeruginosa*. The pathogen *P aeruginosa* (Holloway, 1969) is capable of growth in aerobic or anaerobic conditions [22]. Under aerobic conditions, pyruvate, a product of glycolysis, is transported into the mitochondria where it enters the citric acid cycle. When oxygen is not available, pyruvate undergoes fermentation in the cytoplasm. Instead of being metabolized by cellular respiration, it is converted into waste products. This process allows the regeneration of NAD^++^, which is essential for glycolysis to continue. *P. aeruginosa* uses a pyruvate fermentation pathway to sustain anaerobic survival when alternative anaerobic respiratory and fermentative energy generation systems are absent. This is crucial for the bacterium’s survival in deeper layers of biofilms and in the anaerobic mucus plaques found in cystic fibrosis lungs. Pyruvate fermentation alone cannot sustain significant anaerobic growth of *P. aeruginosa*, but it does provide the bacterium with the metabolic capacity for long-term survival [24].

**Figure 2:**
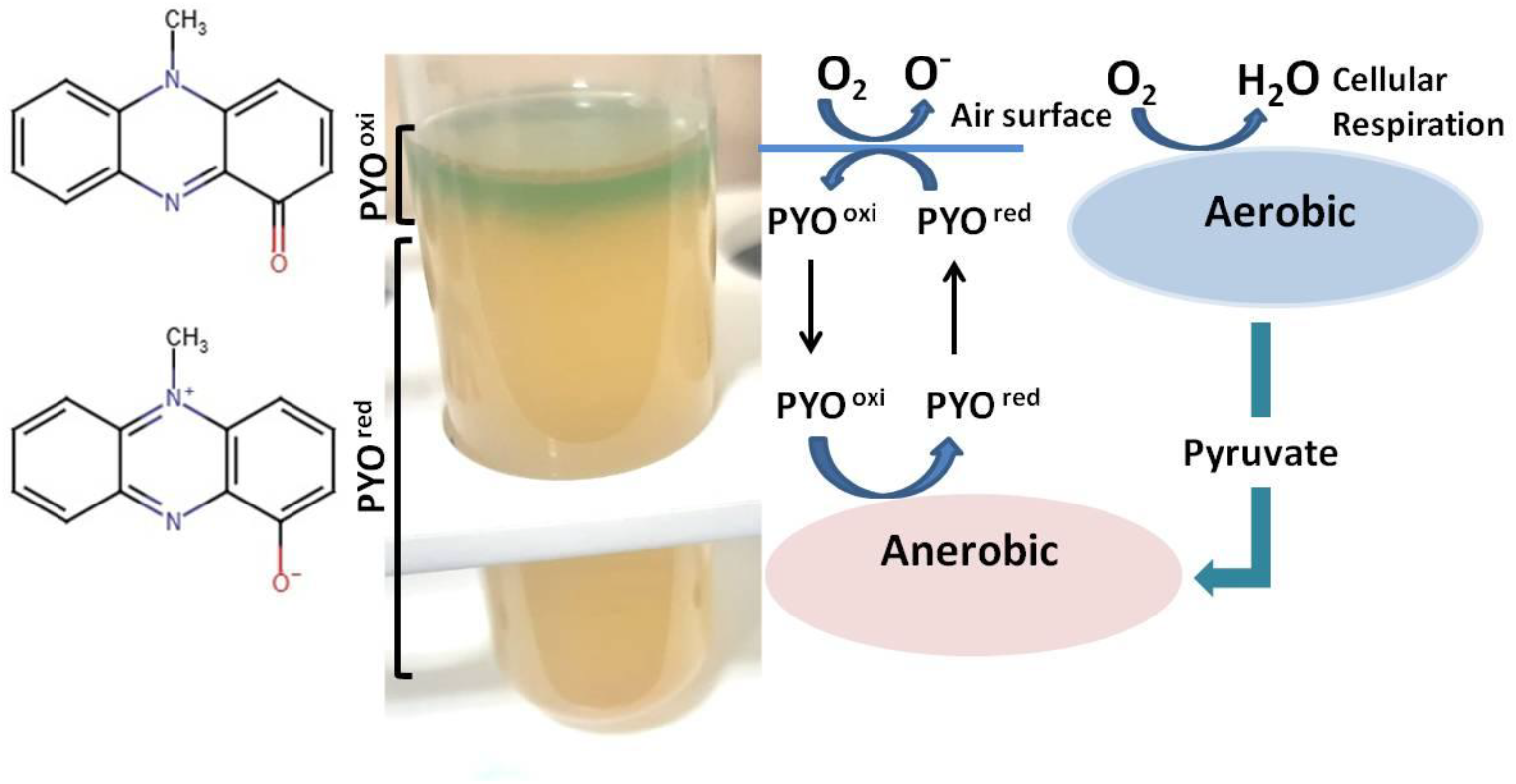
Standing cultures exhibit a green-yellow gradient at the air-medium interface due to pyocyanin (PYO), a redox-active molecule produced by P. aeruginosa. Pyocyanin (PYO) plays a critical role in electron transfer across both aerobic and anaerobic environments, supporting *P. aeruginosa*’s survival in varying oxygen levels. Under anaerobic conditions, PYO cycles between its oxidized (PYO^oxi^) and reduced (PYO^red^) states by accepting electrons from pyruvate, acting as an external electron acceptor. This electron transfer allows bacteria to offload electrons when oxygen is scarce, enabling continued metabolic activity. The reduced pyocyanin then diffuses toward the surface where it can be re-oxidized. In the aerobic region, where cellular respiration occurs, NADH generated from carbon source metabolism donates electrons to the oxidized pyocyanin (PYO^oxi^), reducing it to PYO^red^. This reduced pyocyanin functions as a mobile electron carrier, diffusing to areas where it can be re-oxidized, such as at the air surface. This continuous cycle helps the bacteria maintain redox balance, even in fluctuating oxygen conditions. Overall, pyocyanin acts as a redox mediator, enabling *P. aeruginosa* to survive and adapt by facilitating electron transfer in both oxygen-rich and oxygen-deprived zones.

### 3. UV-Visible Spectroscopy of Pyocyanin

The pyocyanin supernatant underwent UV-visible spectroscopy, which showed a characteristic absorbance peak in the range of 300-400 nm. Upon the addition of hydrochloric acid (HCl) to the pyocyanin solution, a notable color change occurred, shifting from green-blue to pink-red. In the UV-visible spectrum of the pink-red pyocyanin, the peak in the 300-400 nm range was not observed. This colorimetric shift and observed peak change in UV spectrophotometry is attributed to protonation, resulting in a structural rearrangement of the molecule. The protonation of nitrogen atoms within the phenazine ring of pyocyanin plays a crucial role in its chemical behavior and biological functions. Pyocyanin exists in two forms: a zwitterionic state (PYO) and a protonated state (PYOH+), with its form shifting based on the surrounding pH levels [7]. This transition is marked by a distinct colorimetric change, driven by the protonation process that alters the molecular structure and modifies the absorption spectrum of the compound. Such pH-dependent characteristics not only highlight the dynamic nature of pyocyanin but also suggest its potential as an effective colorimetric indicator for monitoring environmental pH changes. This capability could facilitate simple and cost-effective real-time detection of pH fluctuations, making pyocyanin a valuable tool for biosensing applications [12][25].

**Figure 3:**
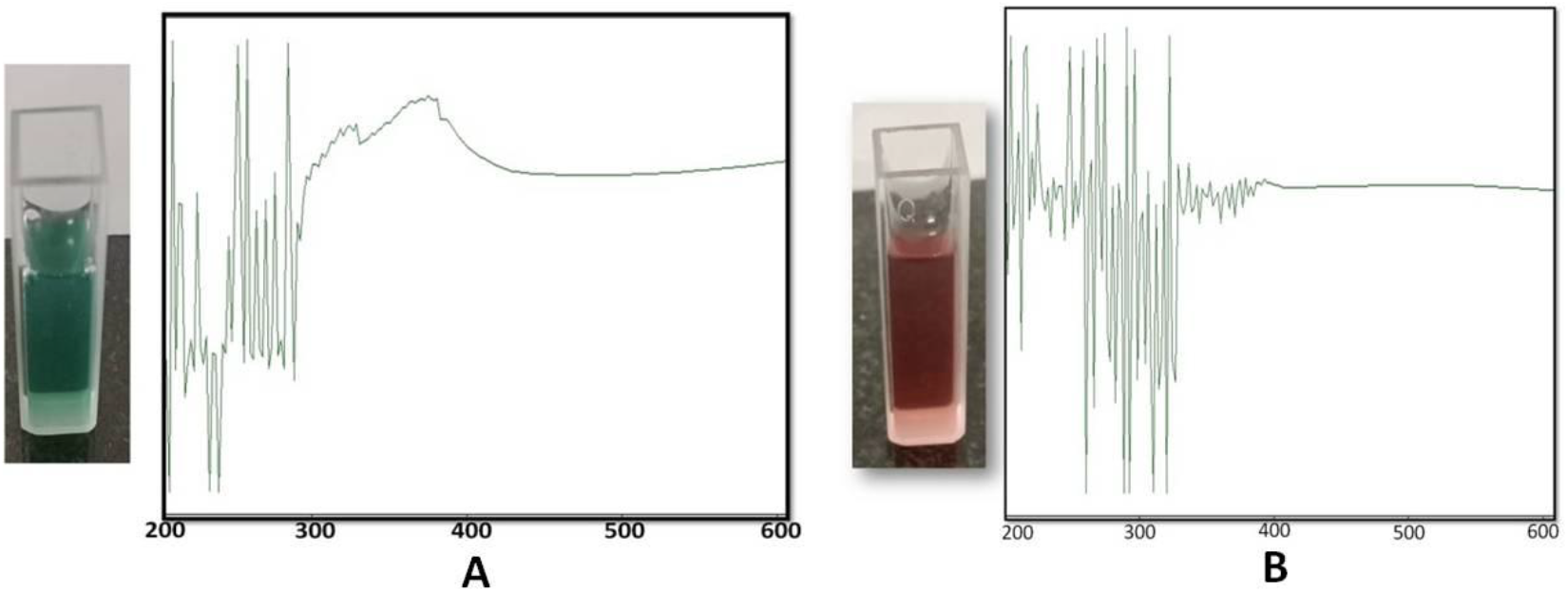
UV absorption spectra of blue colored Pyocyanin (A) and Pyocyanin added with with Hydrochloric acid, red colored (B). The observed colorimetric shift and peak change in UV spectrophotometry are attributed to the protonation-induced structural rearrangement of pyocyanin. Protonation of nitrogen atoms within the phenazine ring governs its chemical properties and biological activity. Pyocyanin exists in a zwitterionic form (PYO) and a protonated form (PYOH+), with its equilibrium state dependent on pH. This transition, characterized by distinct spectral and colorimetric changes, reflects the pH-responsive nature of the molecule.

### 4. Electrochemical Properties of Pyocyanin

The redox properties of pyocyanin (PYO) were examined using cyclic voltammetry. The electrochemical behavior of pyocyanin (culture supernatant) was investigated under different pH conditions using cyclic voltammetry. The protonation of pyocyanin was assessed in both acidic and alkali conditions. The addition of strong acid (HCL) and strong alkali (NaOH) altered the pH levels that influence the protonation process of pyocyanin.

Initially, 1 ml of supernatant pyocyanin solutions (blue in color) were taken in separate tubes, and pH was lowered in respective tubes by adding varying amounts of HCl (2.5 µl, 5 µl, 7.5 µl, and 10 µl), turning to red colour in increasing pH. Electrochemical measurements were conducted for each solution. Additionally, NaOH was added in the lowest pH sample (sample with 10 µl HCL addition) to raise the pH, resulting in a colour change back to the blue region, followed by further electrochemical assessments.

Under acidic conditions, pyocyanin exhibited well-defined reduction of peaks, signifying its ability to protonation. The protonation of the phenazine ring enhances the electrophilicity of the molecule, facilitating electron transfer and positioning pyocyanin as a suitable candidate for redox reactions in acidic environments. In contrast, pyocyanin retained significant redox activity in alkaline conditions, with reduction peaks indicative of stable electron transfer. The dual functionality of pyocyanin across a wide pH spectrum positions it as a versatile redox catalyst [26].

**Figure 4:**
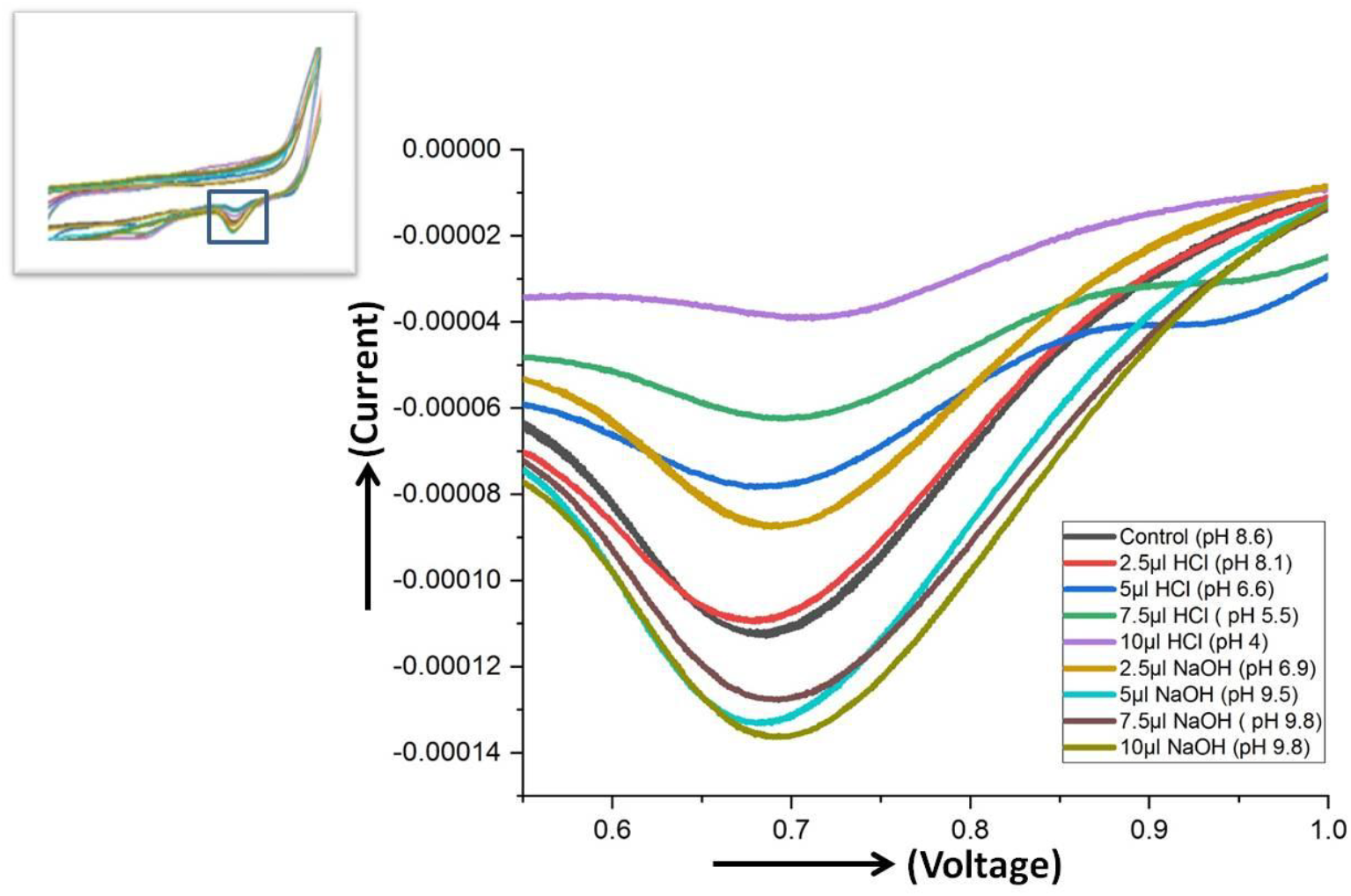
The cyclic voltammetry (CV) responses of an electrochemical system under varying pH conditions, ranging from pH 4 to pH 8.6. The graph plots the current response (A) on the y-axis against the applied voltage (V) on the x-axis, with distinct curves corresponding to each pH level. The CV curves provide insight into how pH influences the redox behavior and electrochemical activity of the system.

### 5. Antimicrobial Properties of Pyocyanin

To perform the antibacterial effect of PYO, we isolated gram positive bacteria from air using the plate exposure method, also known as the settle plate method. We expose LB agar plate in air for 1 hr, after exposure incubates the plates at 37°C for bacteria, and 30°C for fungi overnight. After incubation, some colony was picked up and liquid saturated culture was prepared to test antibacterial property of the PYO. In the same way, agar plate was exposed in air for 6 hrs to get some air fungus. The plate exposure method or passive air sampling is a key technique in environmental and industrial microbiology because it is a cost-effective and simple procedure for monitoring airborne microbial contamination [27]. The principle behind it is the physical settling of airborne particles onto a nutrient agar surface in a Petri dish1. Microorganisms settle on the agar surface, find nutrients and moisture, and grow to form visible colonies.

Previously, we found that three different states of pyocyanin can exist: oxidized (blue), monovalently reduced (colourless) or divalently reduced (red). The conversion of blue to red is done by addition of strong HCL. In our experiment, we decreased gradually the pH of the supernatants, turning towards red colour. The supernatants of different pH were prepared as 8.6, 7.4, 6.6, 5.5, 4.5 and 2.

The antimicrobial efficacy of pyocyanin was assessed at the varying pH ranging alkaline to acidic to determine the impact of pH on its antibacterial potency. Paper disks impregnated with pyocyanin solutions at these pH values were placed on agar plates inoculated with mixed bacterial colonies, and the zones of inhibition were measured.

Zones of inhibition were observed around disks at pH levels above 4.5, with the most significant zones at pH 6.6–7.4. However, at extremely low pH (pH 2 and 4.5), no inhibition zones were formed. This is likely due to excessive protonation at low pH, which decreases the molecule’s ability to effectively interact with bacterial membranes or disrupt metabolic processes. Protonation reduces the redox activity of pyocyanin, limiting its ability to induce oxidative stress in bacterial cells [14].

Conversely, under basic conditions, pyocyanin demonstrated significant antimicrobial activity, with inhibition zones increasing in size as the pH rose. This increase in antibacterial efficacy at higher pH values can be attributed to the deprotonation of pyocyanin, which enhances its redox activity and facilitates electron transfer, potentially destabilizing bacterial cell membranes and interfering with their metabolic processes [12].

The results indicate that pyocyanin’s antimicrobial activity is highly pH-dependent, with optimal activity in neutral to slightly alkaline conditions. This property could be leveraged in environmental applications, such as controlling pathogenic bacteria in aquatic systems or soil remediation, where pH can be modulated to maximize efficacy.

**Figure 5:**
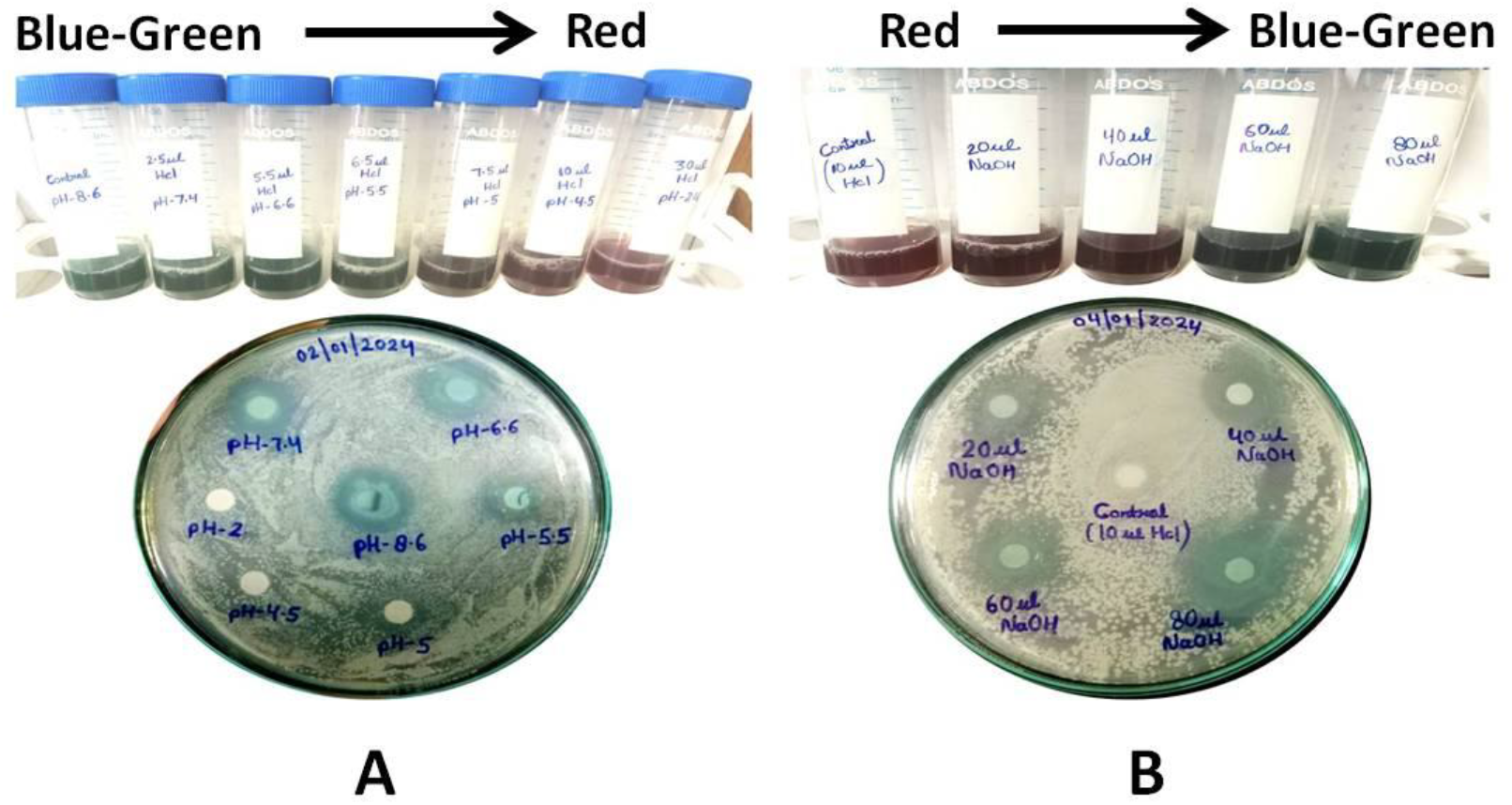
Antimicrobial properties of pyocyanin. Inhibition zones were observed around disks at pH levels above 4.5, with the most pronounced effects at pH 6.6–7.4 (A). In contrast, no inhibition zones formed at extremely low pH (2 and 4.5) (A), likely due to excessive protonation, which reduces pyocyanin’s interaction with bacterial membranes and impairs its metabolic disruption. Protonation diminishes pyocyanin’s redox activity, limiting its ability to induce oxidative stress in bacterial cells. Under alkaline conditions, pyocyanin exhibited enhanced antimicrobial activity, with inhibition zones increasing in size as pH rose (B). This effect is attributed to deprotonation, which enhances redox activity and electron transfer, potentially destabilizing bacterial membranes and interfering with metabolic functions

#### 5.1 Antifungal Activity of Pyocyanin

Pyocyanin’s antifungal properties were similarly assessed using nutrient agar plates inoculated with fungal colonies isolated from soil. Disks impregnated with pyocyanin solutions of varying pH were placed on the plates and incubated at 37°C for 48 hours.

Significant zones of fungal growth inhibition were observed at neutral and basic pH levels, whereas acidic conditions (pH < 5) showed reduced activity. The results revealed significant inhibition of fungal growth at neutral and alkaline pH levels, while acidic conditions exhibited a marked reduction in activity. This diminished activity under acidic conditions is likely attributable to the reduced generation of reactive oxygen species (ROS), which play a crucial role in the oxidative destruction of fungal cells. At neutral and basic pH, pyocyanin likely generates more ROS, disrupting mitochondrial function and electron transport within fungal cells. This dual mechanism of action—generation of ROS and interference with electron transport—positions pyocyanin as a potent antifungal agent in neutral to alkaline environments [12][28].

**Figure 6:**
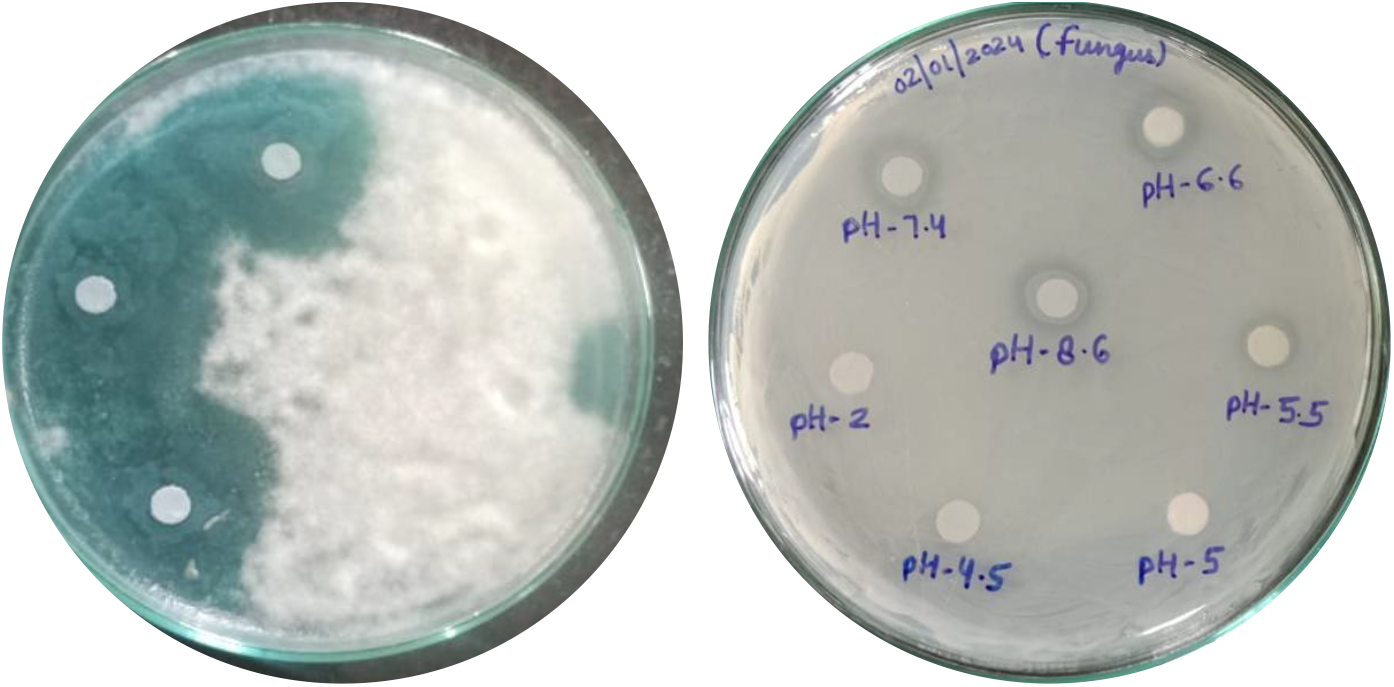
Anti-fungal activity of pyocyanin. Fungal growth inhibition was most pronounced at neutral and alkaline pH levels, while acidic conditions (pH < 5) exhibited significantly reduced activity. This diminished antifungal effect under acidic conditions is likely due to decreased reactive oxygen species (ROS) generation, which is essential for oxidative fungal cell destruction. In contrast, at neutral and basic pH, pyocyanin likely enhances ROS production, disrupting mitochondrial function and electron transport. This dual mechanism-ROS generation and interference with electron transport—highlights pyocyanin’s potency as an antifungal agent in neutral to alkaline environments.

The pH-dependent antimicrobial properties of pyocyanin offer significant implications for medical applications, particularly as a targeted antimicrobial agent. Given that pyocyanin exhibits optimal activity in physiological pH ranges (6.6–7.4), it could be employed in the treatment of infections in human tissues, where the pH is typically neutral to slightly alkaline [12][29]. Additionally, pyocyanin’s redox activity and ability to generate ROS can be harnessed to combat bacterial and fungal infections, particularly in conditions where conventional antibiotics face resistance.

Pyocyanin’s natural origin also makes it an attractive alternative to synthetic antibiotics, aligning with current trends towards the development of environmentally sustainable, naturally derived antimicrobial agents. Given its potential to combat both bacterial and fungal pathogens, pyocyanin could be explored for use in topical treatments for wound infections, as well as in systemic applications for broader-spectrum antimicrobial therapy [16][30].

## CONCLUSION

This study provides a comprehensive evaluation of pyocyanin, a redox-active phenazine derivative produced by *Pseudomonas aeruginosa*, elucidating its electrochemical properties and pH-dependent antimicrobial activities. The results reveal that pyocyanin’s redox functionality and antimicrobial efficacy are closely linked to environmental pH, with its optimal activity observed under neutral to slightly alkaline conditions. The electrochemical analysis through cyclic voltammetry demonstrated that pyocyanin retains its redox activity across a wide pH spectrum, with enhanced reduction peaks in acidic environments and stable electron transfer kinetics in basic conditions. This dual functionality across diverse pH environments highlights pyocyanin’s robustness and adaptability for potential applications in various fields.

The pH-dependent antimicrobial assays confirmed pyocyanin’s strong inhibitory effects on bacterial and fungal pathogens, with the most pronounced activity occurring at pH levels compatible with physiological and environmental conditions. These findings underscore pyocyanin’s promise as an antimicrobial agent, particularly for combating infections caused by multidrug-resistant microorganisms, such as *Staphylococcus aureus* and *Candida albicans*. The ability of pyocyanin to generate reactive oxygen species (ROS) further enhances its antimicrobial properties, positioning it as a potential candidate for use in infection control and therapeutic interventions. However, reduced efficacy under highly acidic conditions indicates the need for tailored formulations to maximize its effectiveness in different environments.

In addition to its antimicrobial properties, the study emphasizes pyocyanin’s potential for environmental and industrial applications. Its redox activity suggests utility in bioremediation and wastewater treatment, particularly in the degradation of synthetic dyes and other recalcitrant pollutants. Moreover, its role as an electron mediator in microbial fuel cells (MFCs) positions pyocyanin as a valuable component in sustainable energy production systems. Furthermore, its structural responsiveness to pH presents opportunities for its use as a biosensing agent in environmental monitoring.

Despite its promising properties, several challenges must be addressed to unlock the full potential of pyocyanin. The cost-effectiveness and scalability of its production remain significant barriers, particularly when considering its application in industrial and clinical settings. Exploring alternative production strategies, such as leveraging marine-derived strains of *P. aeruginosa* or genetically engineered pathways, could significantly improve yields and reduce production costs. Additionally, concerns regarding its stability, potential toxicity, and environmental impact require further investigation to ensure its safe and sustainable application.

In conclusion, this study advances our understanding of pyocyanin’s physicochemical and biological properties, providing valuable insights into its mechanisms of action and versatility. Its dual role as a redox mediator and antimicrobial agent offers a wide range of applications across the medical, environmental, and industrial sectors. Future research should focus on optimizing production methods, improving its functional stability, and exploring synergistic applications to fully harness the biotechnological potential of this unique phenazine compound.

## Author contribution

Mansi Saini Methodology, Investigation, Formal analysis, Writing, Subrata Kumar Das: Conceptualization, Supervision, Methodology, Investigation, Formal analysis, Data curation,Visualization,Validation,Writing, review & editing,. Dharmesh Methodology, Investigation, Gaurav Dutt Investigation, review & editing Kishlay Kant Singh Investigation, review & editing.

